# Does alpha-tocopherol flip-flop help to protect membranes against oxidation?

**DOI:** 10.1101/422006

**Authors:** Phansiri Boonnoy, Mikko Karttunen, Jirasak Wong-ekkabut

## Abstract

Alpha-tocopherols (α-toc) are crucial in protecting biological membranes against oxidation by free radicals. We investigate the behavior of α-toc molecules in lipid bilayers containing oxidized lipids by molecular dynamics (MD) simulations. To verify the approach, the location and orientation of α-toc are first shown to be in agreement with previous experimental results. The simulations further show that α-toc molecules stay inside the lipid bilayer with their hydroxyl groups in contact with the bilayer surface. Interestingly, interbilayer α-toc flip-flop was observed in both oxidized and non-oxidized bilayers with significantly higher frequency in aldehyde lipid bilayer. Free energy calculations were performed and estimates of the flip-flop rates across the bilayers were determined. As the main finding, our results show that the presence of oxidized lipids leads to a significant decrease of free energy barriers and that the flip-flop rates depend on the type of oxidized lipid present. Our results suggest that α-toc molecules could potentially act as high efficacy scavengers of free radicals to protect membranes from oxidative attack and help stabilize them under oxidative stress.

## Introduction

An imbalance between the production and elimination of oxidizing lipid species in cell membrane can lead to oxidative attack on unsaturated lipids.^1^ α-tocopherol (α-toc; a form of Vitamin E) is the most abundant and important lipophilic antioxidant found in cell membranes.^2-4^ It is known to act as an essential radical scavenger protecting membranes from oxidation, but several key aspects of the molecular mechanisms of this action remain uncovered. α-toc inhibits lipid peroxidation process by donating a hydrogen atom to a lipid-peroxyl radical and becoming an α-tocopheroxyl radical.^5^ Although chemical assessment of α-toc’s antioxidant ability has been shown, the mechanisms by which it stabilizes membranes are debated.^3,6-9^

Significant experimental and computational effort has been invested in characterizing the location and interactions of α-toc in model membranes ^8-19^ but obtaining precise information has been proven to be challenging,^9,16^ and the behavior and interaction mechanisms of α-toc molecules inside an oxidized lipid bilayer are being debated. Biophysical studies have, however, demonstrated that the chromanol ring of α-toc is located at the lipid-water interface.^12,13,16^ This suggests that membrane surface is the key to α- toc’s antioxidant activity. It has also been suggested that α-toc partitions in membrane regions are rich in polyunsaturated lipids but more studies are needed to confirm this.^8,9^

In addition to the above, MD simulations at elevated temperatures have demonstrated the presence of transmembrane flip-flops.^10,11^ It has been shown that the flip-flop rate depends on temperature, the degree of unsaturated lipid chains, and the fluidity of the lipid membrane.^10,11^ It has been shown that flip-flops are intimately related to trapping free radicals near the lipid-water interface allowing for the lipid peroxyl radical(s) to be removed.^16^ However, thus far there is no information on how α-toc behaves after lipid peroxidation has occurred. In addition, the functions and molecular insight into α-toc’s actions in oxidative cell membrane remain unclear. In our previous MD simulations^18^ we have shown that α-toc inhibits pore formation in oxidized lipid bilayers. In particular, α-tocs help to stabilize the bilayer structure by trapping the polar oxidized functional group at the water interface thus reducing water permeability across the bilayer. In addition, transmembrane flip-flop of α-toc were observed in both oxidized and non-oxidized bilayer even at room temperature. However, the energetics of the process need to be elaborated.

In this work, we focus on the dynamics and kinetics of α-toc flip-flop in both pure and oxidized lipid bilayers. The effects of different oxidized functional groups were studied. The free energy profiles for α-toc desorbing out of the bilayer were calculated and the flip-flop rates were estimated. The results show that the free energy barriers become significantly decreased in the presence of oxidized lipids, and that the flip-flop rates depend on the type of oxidized lipid present. This study presents new evidence of how α-toc protects oxidized cell membranes.

## Methodology

We performed MD simulations of oxidized and non-oxidized phospholipid bilayers at various α- toc concentrations. All of the lipid bilayers had 128 lipid molecules (64 per leaflet). A 100% 1-palmitoyl-2-lauroyl-sn-glycero-3-phosphocholine (PLPC) lipid bilayer was used as a reference to characterize the effects of α-toc on a non-oxidized bilayer. For the oxidized bilayers, we used 1:1 binary mixtures of PLPC and its four main oxidative derivative products, namely, two hydroperoxides (1-palmitoyl-2-(9-hydroperoxytrans-10, cis-12-octadecadienoyl)-sn-glycero-3-phosphocholine, 9-tc and 1-palmitoyl-2-(13-hydroperoxy-trans-11,cis-9-octadecadienoyl)-sn-glycero-3-phosphocholine, 13-tc), and two aldehydes (1-palmitoyl-2-(9-oxo-nonanoyl)-sn-glycero-3-phosphocholine, 9-al and 1-stearoyl-2-(12-oxo-cis-9-dodecenoyl)-snglycero-3-phosphocholine, 12-al). The molecular structures of α-toc and the lipids are shown in Figure 1. We studied bilayers with 0, 2, 4, 8 and 16 α-toc molecules (equivalent to the concentrations of 0%, 1.5%, 3.0%, 5.9%, 11.1%, respectively). All systems were solvated in 10,628 simple point charged (SPC) water.^20^ The parameters of both PLPC and the oxidized lipids were taken from previous studies.^21-23^ Detailed descriptions of the topologies and force field parameters of α-toc are provided in Refs.^10,11^. Initially, α-toc molecules were randomly placed in the water phase at about 4.2 nm in the *z*-direction from the center of the bilayer. The passive penetration times of the α-toc molecules into the bilayers varied from tens to several hundreds of nanoseconds. At the high concentrations (5.9% and 11.1%) α-toc molecules required much longer times to enter the lipid bilayer (over several microseconds) and in some cases pore formation occurred before complete translocation of α-toc molecules into the lipid bilayer. To avoid artificial pore formation induced by the initial conditions, the α-toc molecules were randomly inserted at the lipid-water interface at high concentrations. The details of all simulations are provided in Table 1.

**Table 1.** The list of systems studied here. Each 100% PLPC bilayer and 50% peroxide (13-tc, 9-tc) mixture was run for 1 μs. The 50% aldehyde (12-al, 9-al) mixtures were run for 1-5 μs to study the effects of α-toc on pore formation. Pore formation time and when a pore/pores occurred is shown in the brackets in the column marked final structure. Both the absolute number and the percentage of α-toc molecules are given in the column description of the system.

| No. | description of the systems | simulation time (ns) | final structure |
| --- | --- | --- | --- |
| 1 | 100% PLPC | 1000 | Bilayer |
| 2 | 100% PLPC + 2 α-toc (1.5%) | 1000 | Bilayer |
| 3 | 100% PLPC + 4 α-toc (3.0%) | 1000 | Bilayer |
| 4 | 100% PLPC + 8 α-toc (5.9%) | 1000 | Bilayer |
| 5 | 50% 13-tc | 1000 | Bilayer |
| 6 | 50% 13-tc + 2 α-toc (1.5%) | 1000 | Bilayer |
| 7 | 50% 13-tc + 4 α-toc (3.0%) | 1000 | Bilayer |
| 8 | 50% 13-tc + 8 α-toc (5.9%) | 1000 | Bilayer |
| 9 | 50% 9-tc | 1000 | Bilayer |
| 10 | 50% 9-tc + 2 α-toc (1.5%) | 1000 | Bilayer |
| 11 | 50% 9-tc + 4 α-toc (3.0%) | 1000 | Bilayer |
| 12 | 50% 9-tc + 8 α-toc (5.9%) | 1000 | Bilayer |
| 13 | 50% 12-al | 1000 | Bilayer with a pore (576 ns) |
| 14 | 50% 12-al + 2 α-toc (1.5%) | 1500 | Bilayer with a pore (1064 ns) |
| 15 | 50% 12-al + 4 α-toc (3.0%) | 3000 | Bilayer with a pore (2802 ns) |
| 16 | 50% 12-al + 8 α-toc (5.9%) | 4000 | Bilayer with a pore (3394 ns) |
| 17 | 50% 12-al + 16 α-toc (11.1%) | 5000 | Bilayer |
| 18 | 50% 9-al | 1000 | Bilayer with a pore (577 ns) |
| 19 | 50% 9-al + 2 α-toc (1.5%) | 1000 | Bilayer with a pore (391 ns) |
| 20 | 50% 9-al + 4 α-toc (3.0%) | 5000 | Bilayer with a pore (4794 ns) |
| 21 | 50% 9-al + 8 α-toc (5.9%) | 2000 | Bilayer with a pore (1589 ns) |
| 22 | 50% 9-al + 16 α-toc (11.1%) | 5000 | Bilayer |

**Figure 1.**
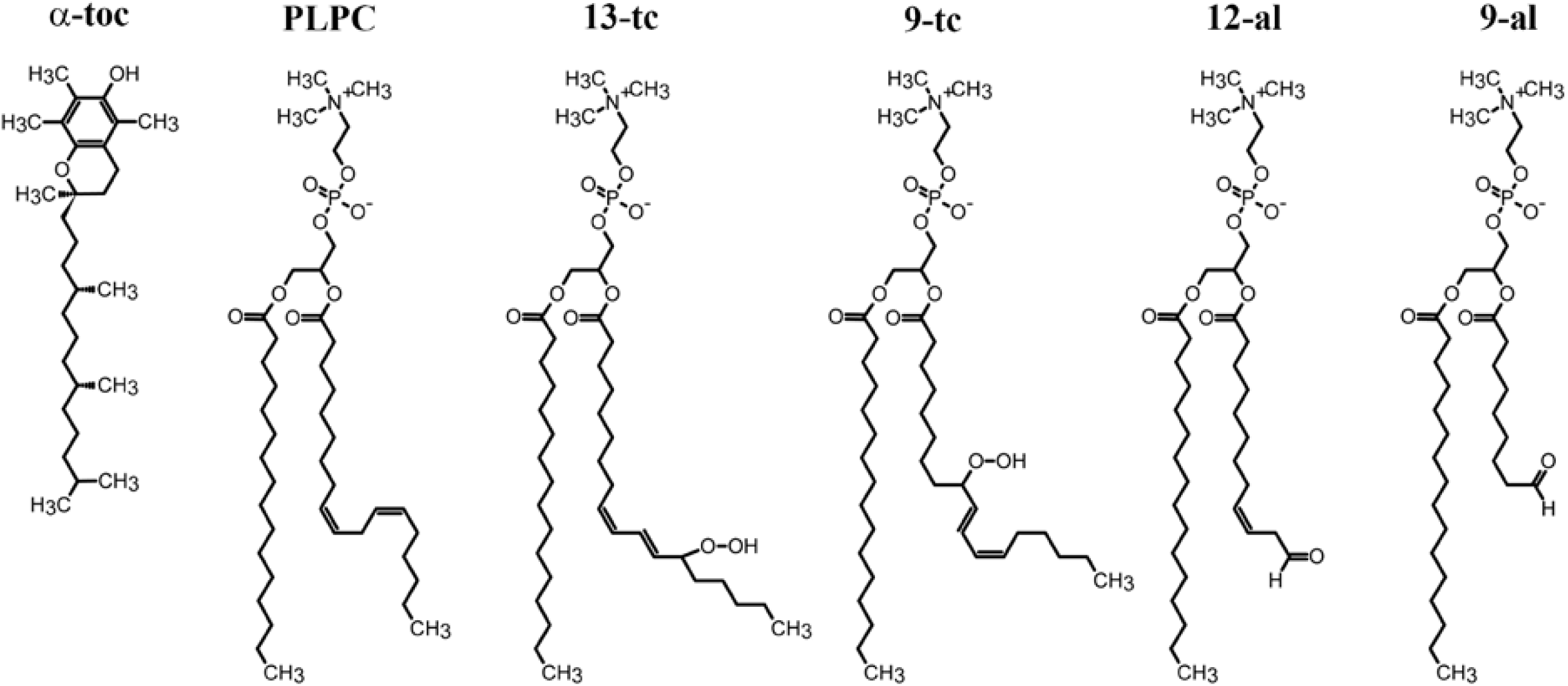
Chemical structures of the alpha-tocopherol (α-toc) and lipid molecules. From left to right: α- toc, 1-palmitoyl-2-lauroyl-sn-glycero-3-phosphocholine (PLPC), 1-palmitoyl-2-(13-hydroperoxy-trans-11,cis-9-octadecadienoyl)-sn-glycero-3-phosphocholine (13-tc), 1-palmitoyl-2-(9-hydroperoxytrans-10, cis-12-octadecadienoyl)-sn-glycero-3-phosphocholine (9-tc), 1-stearoyl-2-(12-oxo-cis-9-dodecenoyl)- snglycero-3-phosphocholine (12-al) and (1-palmitoyl-2-(9-oxo-nonanoyl)-sn-glycero-3-phosphocholine (9-al).

### Simulation Details

After energy minimization using the steeped descents algorithm, MD simulations were run for 1-5 μs with 2 fs integration time step using the GROMACS 5.1.1 package^24^. All simulations were performed in the constant of particle number, pressure and temperature (NPT) ensemble. The v-rescale algorithm^25^ at 298 K with a time constant of 0.1 ps was used for temperature control and equilibrium semi-isotropic pressure was set at 1 bar using the Parrinello–Rahman algorithm^26^ with a time constant of 4.0 ps and compressibility of 4.5 x 10^-5^ bar^-1^. Periodic boundary conditions were applied in all directions and the neighbor list was updated at every time step. A cutoff of 1.0 nm was applied for the real space part of electrostatic interactions and Lennard-Jones interactions. The particle-mesh Ewald method^27-29^ was used to compute the long-range part of electrostatic interactions with a 0.12 nm grid in the reciprocal-space interactions and cubic interpolation of order four. All bond lengths were constrained by the P-LINCS algorithm.^30^ The optimized parameters and protocols have been extensively tested and used previously, for example, in Refs. ^18,23,31-33^. All visualizations were done using Visual Molecular Dynamics (VMD) software.^34^

### Free Energy Calculations

The umbrella sampling technique^35^ with the Weighted Histogram Analysis Method^36^ (WHAM) was used to calculate the potential of mean force (PMF) for transferring an α-toc through the 100% PLPC, 50% 13-tc, 50% 9-tc, 50% 12-al, and 50% 9-al bilayers. For each system, a series of 41 simulation windows was run to compute the PMF profile as a function of the distance between α-toc and the bilayer center varying from z = 0 nm (bilayer center) to z = 4.0 nm (water phase) with 0.1 nm increments. A harmonic potential was applied between the center of mass of the bilayer and the α-toc hydroxyl group with a harmonic force constant of 3000 kJ/(mol nm^2^). Each window was run in the NPT ensemble at 298 K for at least 50 ns; the total time of each PMF profile was at least 2.05 μs. The last 20 ns was used for analysis. The statistical uncertainty in umbrella sampling simulations was estimated by the bootstrap analysis method.^37^

## Result and Discussion

A series of MD simulations of α-toc molecules in different lipid bilayers was performed to study their dynamics in both oxidized and non-oxidized bilayers. Previous MD simulations^18^ have shown that the presence of about 6%-11% of α-toc molecules in an aldehyde bilayer helps protect against passive pore formation induced by oxidized lipids^38^. In this work, pore formation was observed in aldehyde bilayers at 5.9% of α-toc, but the time to form a pore extended for over 1-3 μs. We confirmed that 11.1% α-toc in aldehyde bilayers can prevent pore formation and help to stabilize the oxidized bilayer for times exceeding 5 μs (Figures 2A-B). Interestingly, MD simulations also showed that α-toc’s orientations and transmembrane flip-flops are different in non-oxidized and oxidized bilayer. This indicates that the interactions of α-toc with the bilayer are different before and after lipid peroxidation. It may also indicate, although must be tested by experiments, that these differences are important in determining α-toc’s antioxidant action. To characterize the flip-flops, we calculated the free energy profiles for α-toc desorbing out of the bilayer and estimated the rate of α-toc flip-flop. These issues will be discussed below.

**Figure 2.**
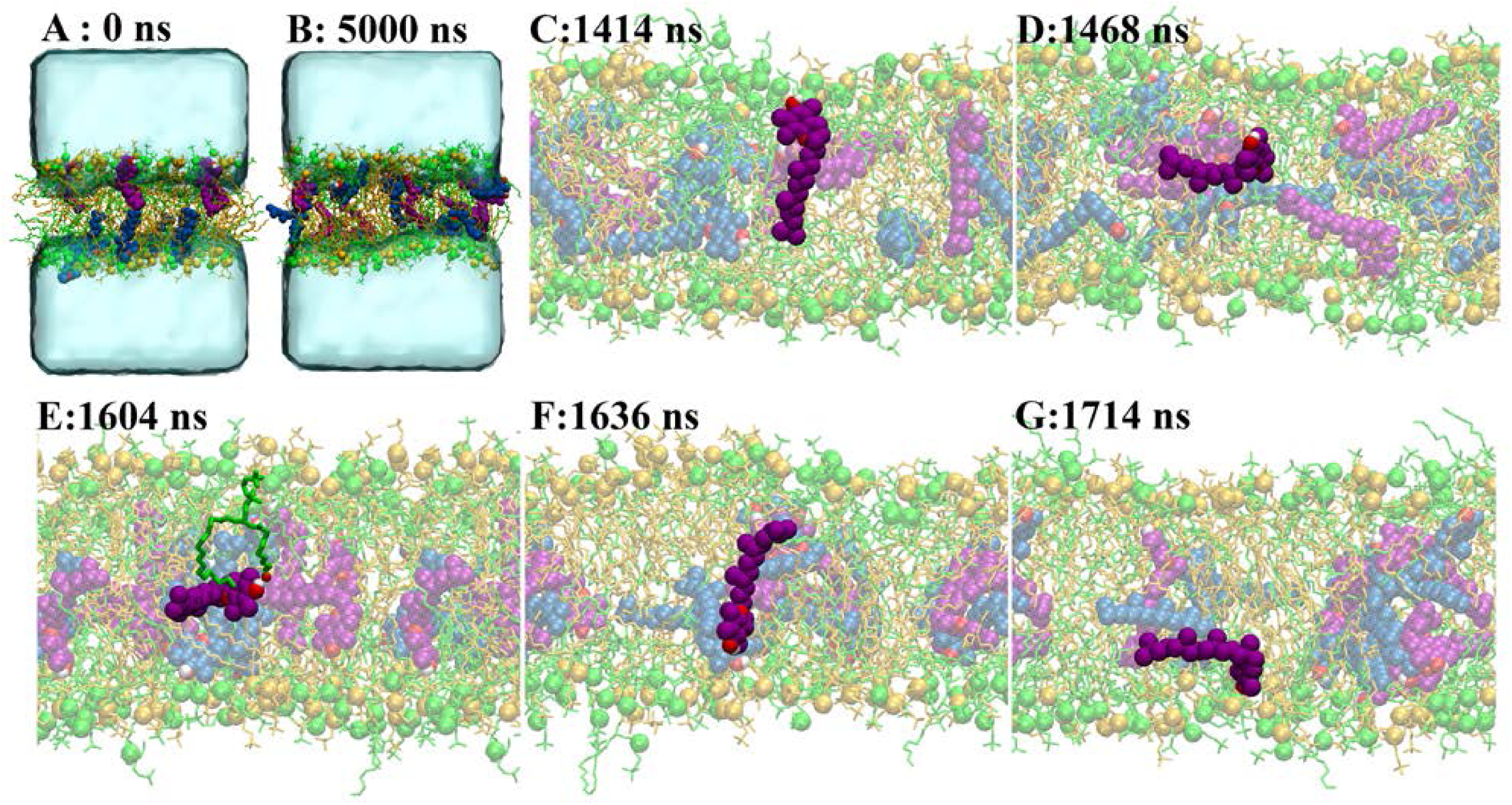
α-toc flip-flop in a 50% 9-al bilayer with 11.1% α-toc molecules present. (A) Initial structure at 0 ns. The 16 α-toc molecules were randomly inserted at the lipid-water interface (8 molecules in each leaflet). PLPC and 9-al lipids are shown as yellow and green lines, respectively, with phosphorus atoms as spheres. α-toc molecules are shown in purple and blue spheres which represent α-tocs in the upper and lower leaflets, respectively. Water is shown in light blue (only in panel A for clarity). Oxygen and hydrogen atoms of the α-toc molecules are shown in red and white spheres, respectively. (B) The final structure at 5 μs (panel A) shows that α-tocs have moved (via flip-flop) between the two leaflets. (C)-(E) The α-toc molecule reorients itself in order to move toward the bilayer center. Within the aldehyde oxidized bilayer, α-toc’s hydroxyl group can form a hydrogen bond with the oxidized lipid tail (E) leading to an increase in the flip-flop rate in the oxidized bilayer. Reorientation of α-toc molecules was observed when they moved to the lower leaflet with the hydroxyl group facing the lipid headgroup and the hydrophobic tail randomly spreading within the bilayer (F-G).

### The behavior of α-toc inside the lipid bilayers

To investigate where α-toc molecules reside inside the different bilayers, we investigated the time evolution of the positions of α-tocs’ hydroxyl groups along the *z*-axis (Figure 3). The results show the hydroxyl groups prefer to stay at the bilayer-water interface around the carbonyl group (see Figure 1 for the lipid structures) for both oxidized and non-oxidized bilayers, Table S1. This is in agreement with previous MD simulations^18^ and experiments^9^.

**Figure 3.**
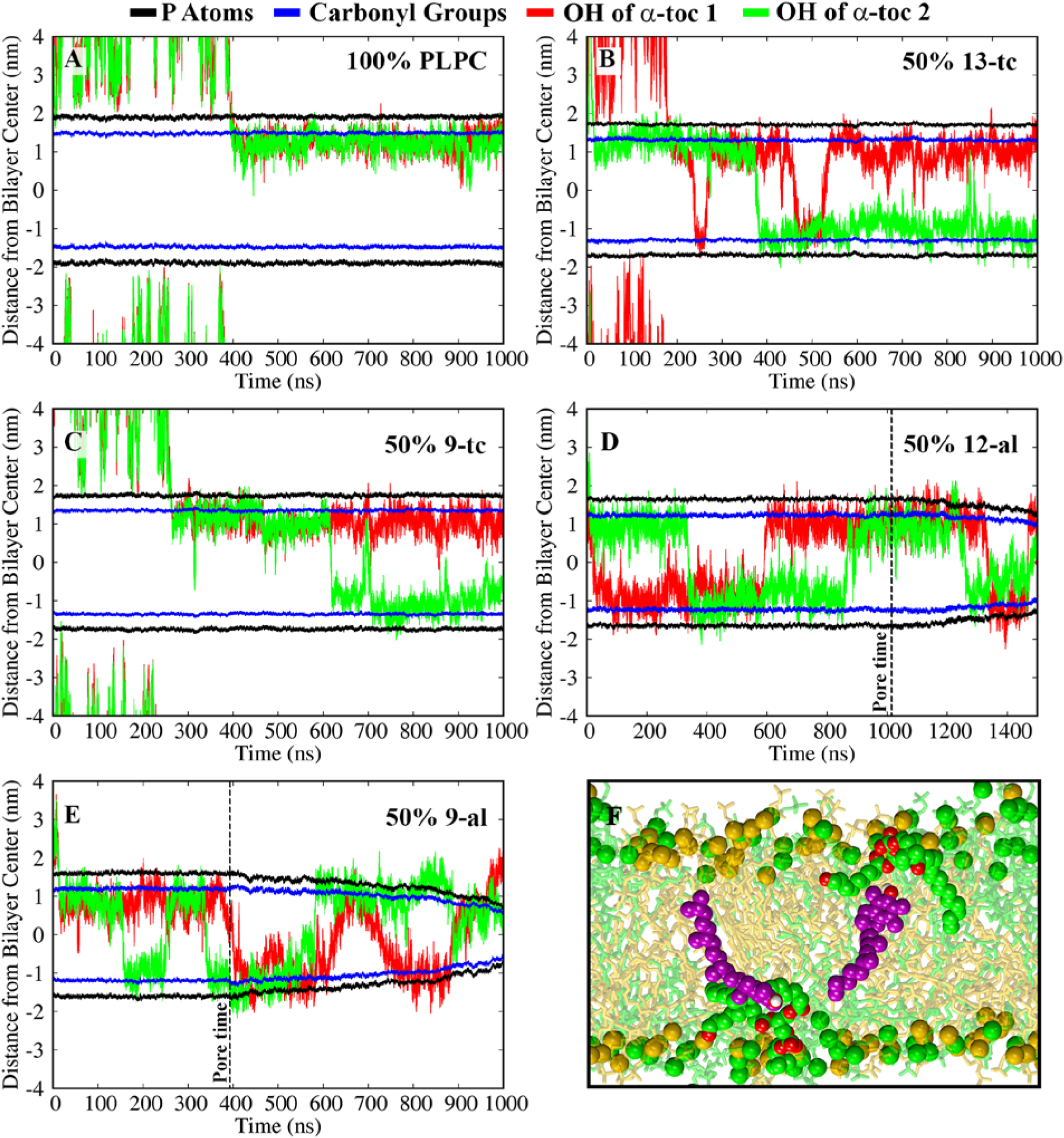
(A-E) Time evolution of the distance of α-toc’s hydroxyl group from the bilayer center along the *z-*axis in non-oxidized and oxidized lipid bilayers containing 2 α-toc molecules. The dashed vertical lines show the times when pore formation first started to occur. Note that the definition of bilayer center becomes somewhat ambiguous during pore formation. (F) The preferred location of α-toc molecules inside the 50% 9-al bilayer is underneath the lipid carbonyl group. The color code for the lines is given on top of the picture.

In non-oxidized membranes, the presence of α-toc molecules in the lipid-water interface region helps to protect the bilayer from approaching free radicals.^5^ When oxidized lipids are present, α-toc flip-flop has an important function in helping to reach and scavenge peroxyl radicals within the lipid bilayer.^16^ Figures 2C-G show snapshots of α-toc transmembrane flip-flop. In the next two sections we quantify the process by computing the free energy and the flip-flop rates, but as an indicative rough comparison we followed the molecules in five lipid bilayers containing 5.9% α-toc molecules over 1 μs and found 21, 10, 16, 7, and 4 events for 50% 9-al, 50% 12-al, 50% 13-tc, 30% 9-tc, and 100% PLPC lipid bilayers, respectively. For the aldehyde bilayers containing 11.1% α-toc molecules, we observed 136 flip-flops for the 50% 9-al bilayer and 162 for the 50% 12-al bilayer over 5 μs. This suggests that the faster dynamics of the α-toc molecules may be a key to its antioxidant action and consequent stabilization of the bilayer under oxidative stress.

The orientations of the α-toc molecules were investigated by calculating the angle between the bilayer normal (*z*-axis) and a vector connecting the last methyl group in the tail to the hydroxyl group in the head group (see Figure 1 for the molecular structures). The last 500 ns (before pore creation in aldehyde bilayers) of lipid bilayers with 5.9% α-toc molecules were used for analysis and shown in Figure 4 and Figure S1. The angles range between 0 and 90 degrees: zero degrees equals to alignment parallel to the bilayer normal and 90 degrees to being perpendicular to the bilayer normal. For the PLPC lipid bilayer, most of the α-toc’s hydroxyl groups were found to be located slightly underneath the lipids’ carbonyl groups within the lipid bilayer (Table S1) in agreement with previous experimental studies.^8,9,19^ The tilt angles of the α-toc molecules with respect to the bilayer normal are shown in Figure 4A. Two possible orientations at around 40±12 degrees and 90±4 degrees to the bilayer normal, were observed in PLPC bilayer. In contrast to experiments, the chains of α-toc and saturated DMPC (1,2-dimyristoyl-sn-glycero-3-phosphocholine) and DPPC (1,2-dipalmitoyl-sn-glycero-3-phosphocholine) lipids are oriented with the relative angles of 11±2 and 16±4 degrees, respectively which may imply to the only parallel orientation to the bilayer normal.^19^ However, the α-toc tilt angle is still being debated since discrepancies can arise from differences in lipid types, lipid phase, and location of α-toc inside the bilayer.^10,17,19^ In an analogous manner, differences in tilt angle have been shown to indicate the ability of different sterols to undergo flip-flop.^39^ For a general discussion of lipid flip-flops, see articles by Gurtovenko and Vattulainen^40^, and Sapay, Bennett and Tieleman^41^.

**Figure 4.**
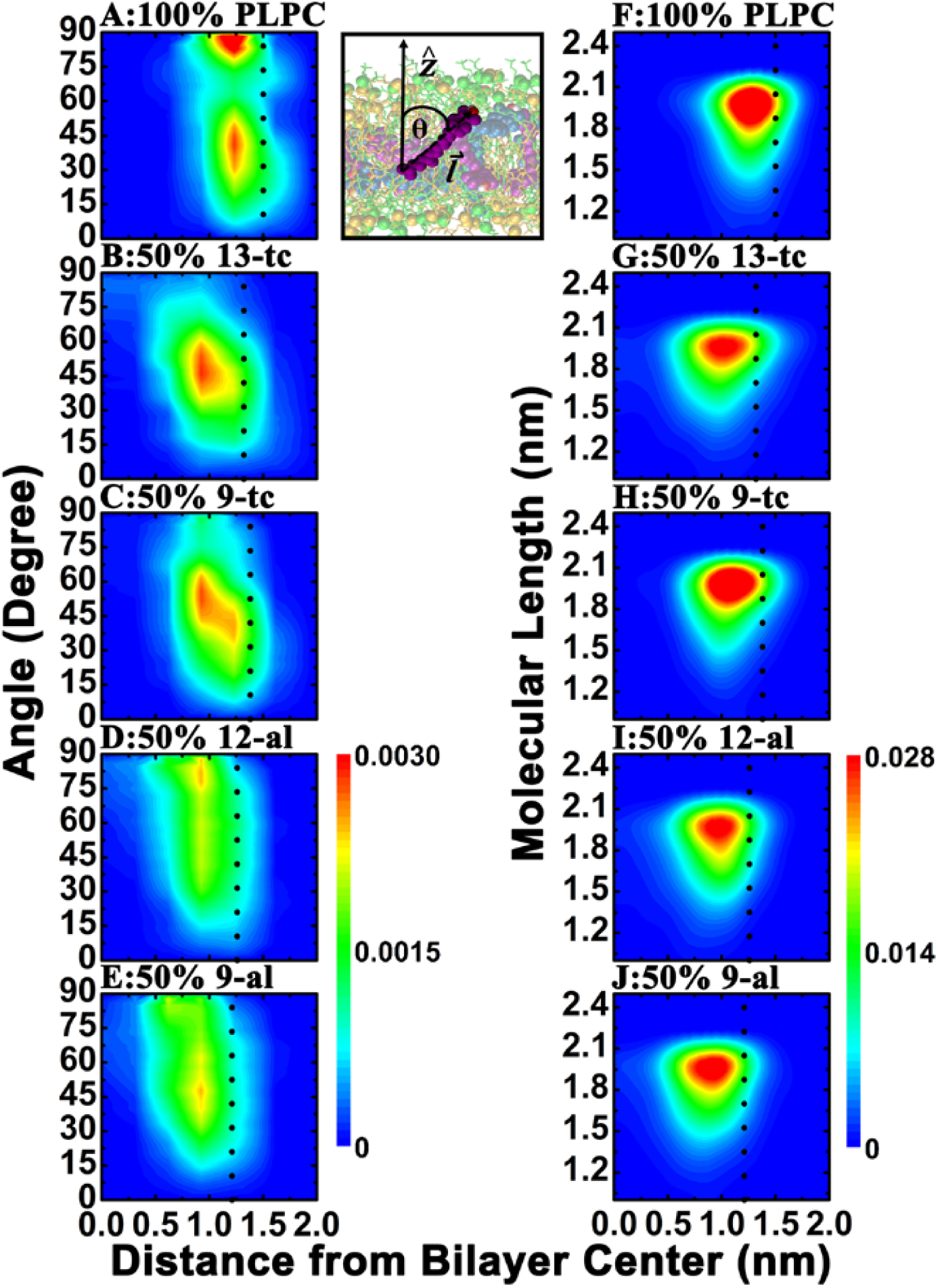
(A-E): Normalized 2D histograms of α-toc’s tilt angle in non-oxidized and oxidized bilayers containing 5.9% α-toc molecules. The hydroxyl group in the chromanol ring and the methyl group at the terminal chain are represented head and tail of α-toc, respectively. The distance from head to tail of α-toc is defined as the molecular length 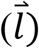 (F-J): α-toc’s molecular length is plotted as a function of the distance of α-toc’s head from the bilayer center. Inset: The tilt angle is defined by the angle between the bilayer normal (*z*-axis) and the vector connecting the last methyl group in the tail to the hydroxyl group in the head of the α-toc molecule. The tilt angle of 0 degrees represents α-toc oriented in parallel to the bilayer normal (*z*-axis) and 90 degrees describes the α-toc being perpendicular to the bilayer normal. The dotted lines represent the average positions of the carbonyl groups in lipid chains.

For the hydroperoxide lipid bilayer, the polar head groups of the α-toc molecules prefer to stay at the bilayer-water interface in a way similar to what is observed for the PLPC bilayer. The average distances of the carbonyl group from the bilayer center are 1.32±0.01 and 1.37±0.02 nm for 50% 13-tc and 50% 9-tc bilayer, respectively. Meanwhile, the locations of the hydroxyl group of α-toc molecules in 50% 13-tc and 50% 9-tc bilayer are 1.03±0.02 and 1.08±0.04 nm from the bilayer center, respectively. However, all α-toc molecules in the peroxide bilayer prefer to be slightly tilted in parallel to the bilayer (Tilted angle is around 40-60 degree to the bilayer normal.) rather than to lie horizontally in the bilayer (Figure 4B-C). For the aldehyde lipid bilayer, a wide range of orientations was observed caused by a random spread of the α-toc’s hydrophobic tail in the bilayer (Figure 4D-E). Interestingly, it can be seen that when the α-toc’s hydroxyl moves deep inside the bilayer and reorients, a flip-flop typically follows (Figure 4D-E). The molecular length 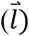 of α-toc in lipid bilayer was determined as the distance between the terminal methyl group in the tail and the hydroxyl group in the head (see Figure 1 for molecular structures). The molecular length distribution of α-toc in non-oxidized/oxidized lipid bilayers is shown in Figure 5. The results show an extended structure in both PLPC and oxidized bilayers with the maximum probability densities in molecular length being approximately 2.06±0.01 nm (Figure 5). The average molecular lengths were found to be 1.88±0.02, 1.87±0.01, 1.91±0.01, 1.85±0.01, 1.86±0.01 nm in PLPC, 13-tc, 9-tc, 12-al, and 9-al lipid bilayers, respectively. Note the emergence of a shoulder in the case of 5.9% 12-al (solid blue curve). As Figure 1 shows, pore formation is observed in this system. These structures are maintained even when the α-toc molecules move toward the bilayer center and undergo a flip-flop to the opposite side of the bilayer (Figure 4F-J). Note the shortened structure of α-toc when they tilt away from bilayer as seen in Figure S2.

**Figure 5.**
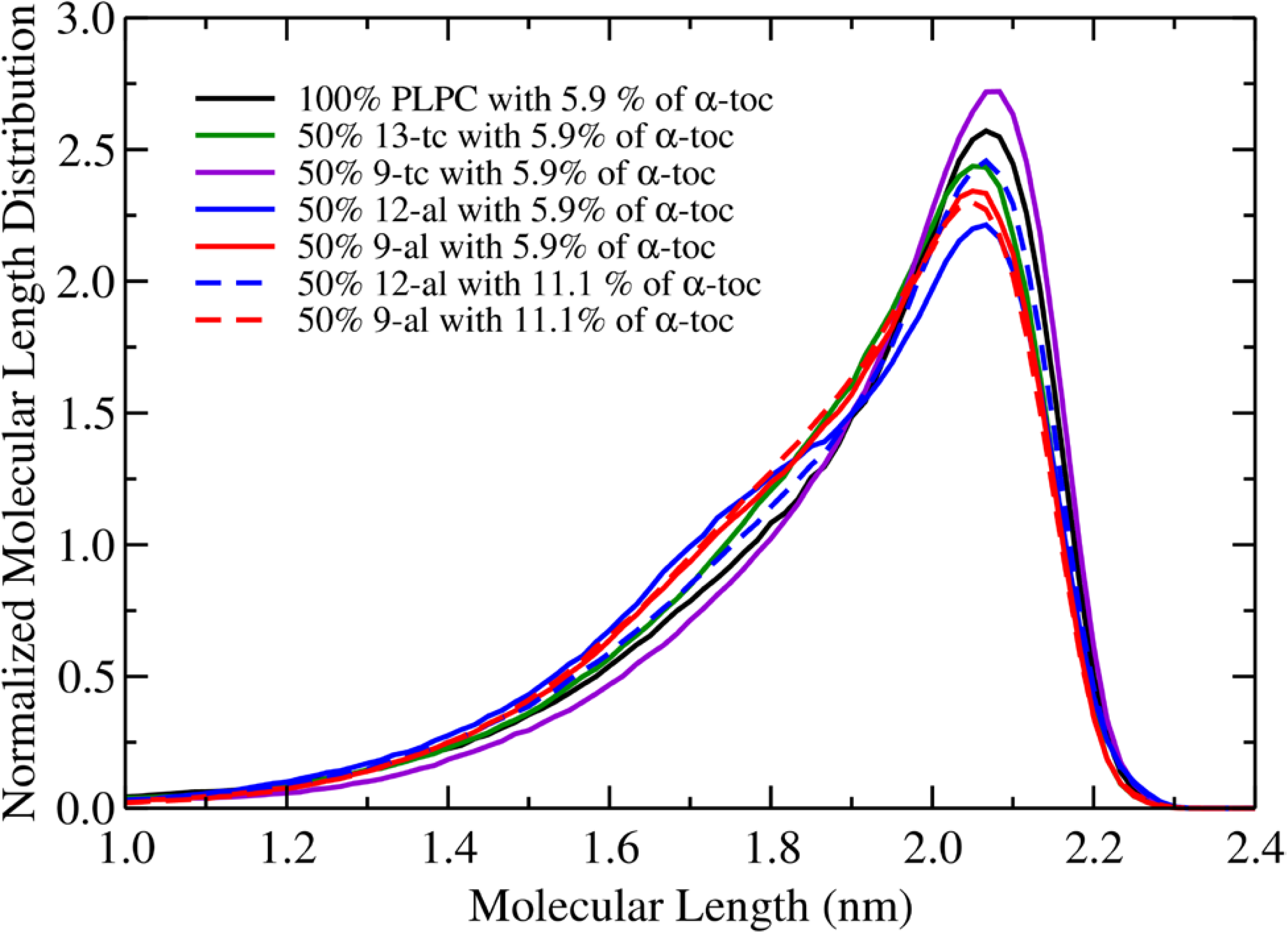
Distribution of α-toc’s molecular length 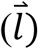 in oxidized and non-oxidized lipid bilayers containing 5.9% and 11.1 % of α-toc molecules. The hydroxyl group in the chromanol ring and methyl group at the terminal chain are represented head and tail of α-toc, respectively. The distance from head to tail of α-toc is defined the molecular length 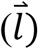.

### Free energy profile of α-toc molecule in the bilayer

Figure 6 shows the potential of mean force (PMF) profiles for moving an α-toc molecule from the bilayer center to the water phase, and Table 2 lists the equilibrium positions (corresponding to the α-toc hydroxyl group), the free energy of desorption (ΔG_desorb_) for moving an α-toc from its equilibrium position in bilayer into bulk water, and the free energy barrier (ΔG_barrier_) for moving an α-toc from the equilibrium position to the bilayer center. Note that thermal energy is about 2 kJ mol^-1^. These results suggest that the probability for an α-toc to undergo flip-flop is higher in oxidized bilayers.

**Table 2.** The equilibrium positions given as the location of the α-toc hydroxyl group from the bilayer center. ΔG_desorb_is the free energy of desorption, that is, the free energy cost for moving an α-toc from its equilibrium position in bilayer into bulk water, and ΔG_barrier_is the free energy for moving an α-toc from its equilibrium position to bilayer center. See also Figure 6.

| systems | $\Delta G_{\text{desorb}}$ (kJ/mol) | $\Delta G_{\text{barrier}}$ (kJ/mol) | equilibrium position (nm) |
| --- | --- | --- | --- |
| 100% PLPC | $-90.0 \pm 0.7$ | $13.5 \pm 0.5$ | $1.26 \pm 0.01$ |
| 50% 13-tc | $-87.8 \pm 1.0$ | $8.1 \pm 0.5$ | $1.10 \pm 0.09$ |
| 50% 9-tc | $-86.6 \pm 0.7$ | $9.2 \pm 0.5$ | $1.24 \pm 0.02$ |
| 50% 12-al | $-87.0 \pm 0.7$ | $8.3 \pm 0.5$ | $1.04 \pm 0.02$ |
| 50% 9-al | $-86.6 \pm 0.5$ | $7.1 \pm 0.2$ | $0.85 \pm 0.01$ |

**Figure 6.**
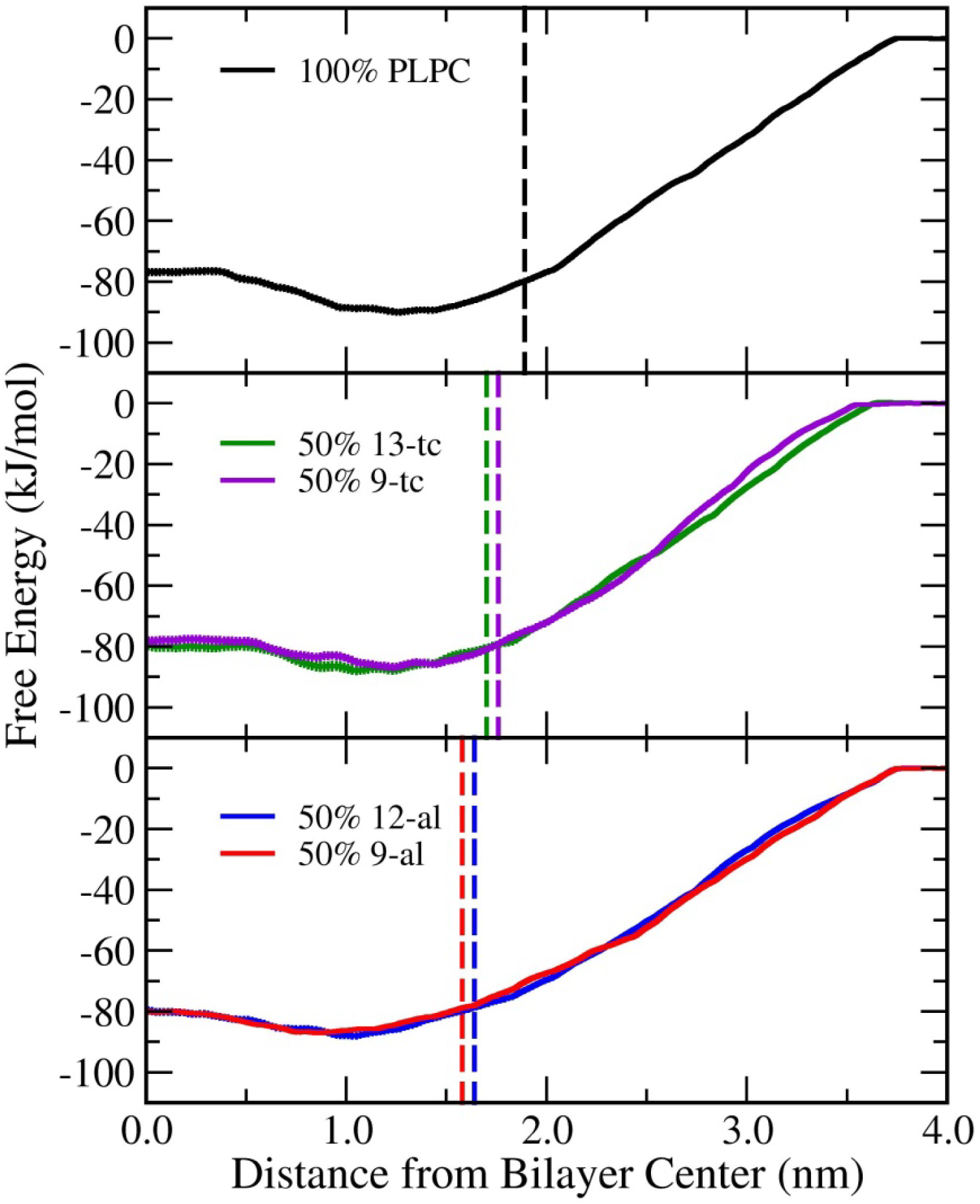
The potential of the mean force (PMF) for moving an α-toc from the center of a 100% PLPC bilayer (A), 50% peroxide lipid (B) and a 50% aldehyde lipid (C) to the water phase as a function of distance in the *z*-direction from the bilayer center. The PMF in bulk water was set to zero. The dashed lines are the average positions of the phosphorus atoms in the lipid head group in each bilayer. The free energy of desorption (ΔG_desorb_) is the difference in free energy between the equilibrium position and bulk water. The free energy barrier (ΔG_barrier_) is the maximum free energy for α-toc to move from the equilibrium position to the bilayer center. All free energy calculations in different lipid bilayers are listed in Table 2. The error in all systems is lesser than 1 kJ mol^-1^.

### Flip-flop rate calculation

To estimate the rate of α-toc flip-flops (*k*_*flip*_), additional simulations with an α-toc initially at the bilayer center were performed. The initial configurations were taken from the *z*=0 nm window in umbrella sampling simulations. 50 independent simulations of each bilayer were independently run for 50 ns, and the time for the hydroxyl group of the α-toc to return from the bilayer center to the equilibrium position (*t*_*d*_) was recorded. The rate of α-toc flip-flop across a lipid bilayer (*k*_*flip*_) was calculated following the approach of Bennett et al.^42^ using the equation

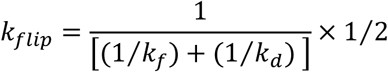

Where

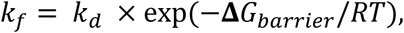

and where *R* and *T* are the gas constant and temperature, respectively. *ΔG*_*barrier*_ is the free energy difference between the equilibrium position and the bilayer center. The half time for flip-flop (*t*_*1*/*2*_) was calculated using^43^

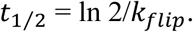

Table 3 shows the results and the parameters. The results show that α-toc molecules undergo rapid flip-flops especially in the aldehyde bilayers as compared to flip-flops in a pure phospholipid bilayer.

**Table 3.**
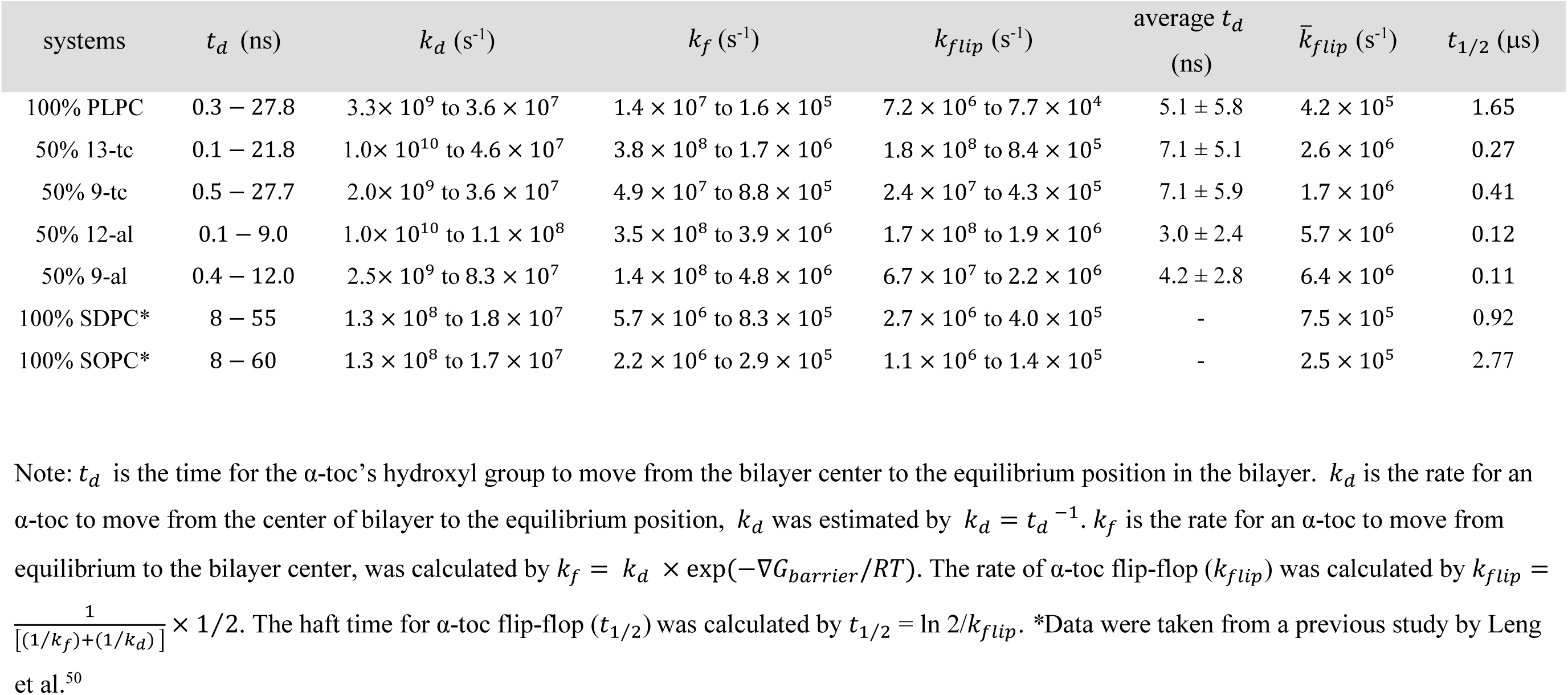
The parameters associated with α-toc flip-flop calculation in different lipid bilayers.

| systems | $t_d$ (ns) | $k_d$ (s<sup>-1</sup>) | $k_f$ (s<sup>-1</sup>) | $k_{flip}$ (s<sup>-1</sup>) | average $t_d$<br>(ns) | $\bar{k}_{flip}$ (s<sup>-1</sup>) | $t_{1/2}$ (μs) |
| --- | --- | --- | --- | --- | --- | --- | --- |
| 100% PLPC | 0.3 – 27.8 | $3.3 \times 10^9$ to $3.6 \times 10^7$ | $1.4 \times 10^7$ to $1.6 \times 10^5$ | $7.2 \times 10^6$ to $7.7 \times 10^4$ | $5.1 \pm 5.8$ | $4.2 \times 10^5$ | 1.65 |
| 50% 13-tc | 0.1 – 21.8 | $1.0 \times 10^{10}$ to $4.6 \times 10^7$ | $3.8 \times 10^8$ to $1.7 \times 10^6$ | $1.8 \times 10^8$ to $8.4 \times 10^5$ | $7.1 \pm 5.1$ | $2.6 \times 10^6$ | 0.27 |
| 50% 9-tc | 0.5 – 27.7 | $2.0 \times 10^9$ to $3.6 \times 10^7$ | $4.9 \times 10^7$ to $8.8 \times 10^5$ | $2.4 \times 10^7$ to $4.3 \times 10^5$ | $7.1 \pm 5.9$ | $1.7 \times 10^6$ | 0.41 |
| 50% 12-al | 0.1 – 9.0 | $1.0 \times 10^{10}$ to $1.1 \times 10^8$ | $3.5 \times 10^8$ to $3.9 \times 10^6$ | $1.7 \times 10^8$ to $1.9 \times 10^6$ | $3.0 \pm 2.4$ | $5.7 \times 10^6$ | 0.12 |
| 50% 9-al | 0.4 – 12.0 | $2.5 \times 10^9$ to $8.3 \times 10^7$ | $1.4 \times 10^8$ to $4.8 \times 10^6$ | $6.7 \times 10^7$ to $2.2 \times 10^6$ | $4.2 \pm 2.8$ | $6.4 \times 10^6$ | 0.11 |
| 100% SDPC* | 8 – 55 | $1.3 \times 10^8$ to $1.8 \times 10^7$ | $5.7 \times 10^6$ to $8.3 \times 10^5$ | $2.7 \times 10^6$ to $4.0 \times 10^5$ | - | $7.5 \times 10^5$ | 0.92 |
| 100% SOPC* | 8 – 60 | $1.3 \times 10^8$ to $1.7 \times 10^7$ | $2.2 \times 10^6$ to $2.9 \times 10^5$ | $1.1 \times 10^6$ to $1.4 \times 10^5$ | - | $2.5 \times 10^5$ | 2.77 |

Note: $t_d$ is the time for the α-toc's hydroxyl group to move from the bilayer center to the equilibrium position in the bilayer. $k_d$ is the rate for an α-toc to move from the center of bilayer to the equilibrium position, $k_d$ was estimated by $k_d = t_d^{-1}$ . $k_f$ is the rate for an α-toc to move from equilibrium to the bilayer center, was calculated by $k_f = k_d \times \exp(-\nabla G_{barrier}/RT)$ . The rate of α-toc flip-flop ( $k_{flip}$ ) was calculated by $k_{flip} = \frac{1}{[(1/k_f) + (1/k_d)]} \times 1/2$ . The half time for α-toc flip-flop ( $t_{1/2}$ ) was calculated by $t_{1/2} = \ln 2/k_{flip}$ . \*Data were taken from a previous study by Leng et al.<sup>50</sup>

To put the results in Table 3 into context, we compare the α-toc flip-flop rates to those reported in different systems. In pure phospholipid bilayers, previous experimental studies have reported *t*_*1*_/_*2*_ in a large unilamellar DPPC vesicle in the fluid phase (temperature range of 50-65 °C) to be in range from days to weeks, and no flip-flop was observed in the gel phase over 250 h.^44^ Previous free energy calculations^45^using umbrella sampling technique have suggested that the free energy barrier for phospholipid flip-flop decreases upon increasing degree of oxidation in 1-palmitoyl-2-oleoyl-glycero-3-phosphocholine (POPC) bilayers which indicates an increase in the lipid flip-flop rate. However, the flip-flop rates were not explicitly determined and the free energy barriers (between bilayer center and equilibrium position) remain relatively high, 65±6 kJ/mol in the case of a 50% peroxide bilayer.^45^ Finally, in previous simulations, the phospholipid flip-flop rates in pure DPPC bilayers have been estimated to occur in timescales of 4-30 h.^46^

Next, we compare the α-toc flip-flop rates with those of cholesterol in different bilayers; It has been shown that cholesterol helps to protect membranes against oxidized lipids.^47,48^ The flip-flop rates of cholesterol in pure DPPC bilayers are range from 1.2×10^4^to 6.6×10^5^s^-1^.^42^ In unsaturated diarachidonyl-PC (DAPC, 20:4-20:4 PC) lipid bilayer, the flip-flop rates (5.2×10^5^ to 3.7×10^6^ s^-1^) were observed to be an order of magnitude larger than in fully saturated DPPC lipid bilayers.^42,49^ Our calculations show (Table 3) that the α-toc flip-flop rate and the flip-flop half time in 100% PLPC bilayer are 4.2×10^5^ s^-1^ and 1.65 μs, respectively. These values are consistent with a previous computational study using SDPC (1-stearoyl-2-docosahexaenoylphosphatidylcholine, 18:0-22:6PC) and SOPC (1-stearoyl-2-oleoylphosphatidylcholine, 18:0-18:1PC) bilayers.^16,50^ From the binding free energy calculations of α-toc with polyunsaturated phospholipids, the flip-flop rates have been estimated to be 7.5×10^5^ s^-1^ and 2.5×10^5^ s^-1^ in SDPC and SOPC lipid, respectively.^50^ Here, for the oxidized lipid systems the flip-flop rates were found to be (all at 50%): 13-tc: 2.6 ×10^6^ s^-1^, 9-tc: 1.7×10^6^ s^-1^, 12-al: 5.7×10^6^s^-1^ and 9-al: 6.4×10^6^s^-1^. Thus, the flip-flop rates of α-toc in the oxidized bilayers are up to an order of magnitude larger than in the pure PLPC bilayer. This result suggests faster α-toc dynamics in oxidized lipid bilayer, thus providing a simple and robust mechanism to increase interactions with free radicals in membranes with oxidized lipids.

## Conclusion

We have carried out a series of MD simulations of α-toc in different oxidized and non-oxidized lipid bilayers to better understand the role of α-toc in oxidative stress. The presence of about 11% of α- toc was confirmed to stabilize oxidized membranes and prevent water pore formation in a long simulation times over 5 μs. We investigated the locations and conformations of α-toc both in oxidized and non-oxidized bilayers. The results show that α-toc’s hydroxyl group favors staying at the bilayer interface for both cases. In addition, α-toc was observed flip-flopping in both oxidized and non-oxidized lipid bilayers. We calculated the free energy profiles to obtain the free energy barrier and to estimate the α-toc flip-flop rate in the different bilayers. Importantly, our results show that the free energy barriers for α-toc flip-flop become significantly suppressed in the presence of oxidized lipids. As a consequence, the flip-flop rate increases with lipid peroxidation by up to an order of magnitude compared to the pure PLPC bilayer. The rate increases in the following order: aldehyde > peroxide > PLPC. This significantly increased flip-flop rate provides a physical mechanism which allows α-toc to scavenge radicals in order to protect membranes from oxidative attack and to help stabilize membrane under oxidative stress.

Our results indicate that α-toc flip-flop may indeed be essential for membrane protection. To elaborate its role, it would be interesting to investigate its kinetics in multicomponent membranes containing polyunsaturated lipids and in the presence of cholesterol. Such studies would help answer questions about synergistic effects of cholesterol and α-toc, possible aggregation behaviors, and the locations, both vertical and lateral, of both oxidized lipids and α-toc.

## Acknowledgements

This work was financially supported by Kasetsart University Research and Development Institute (KURDI) and Faculty of Science at Kasetsart University (JW). The support of the Thailand Research Fund (TRF) through the Royal Golden Jubilee Ph.D. Program (Grant No. PHD/0204/2559) and the TRF Research Scholar Program (Grant No. RSA6180021) for PB and JW, respectively is acknowledged. MK would like to thank the Natural Sciences and Engineering Research Council of Canada (NSERC) and the Canada Research Chairs Program. Computing facilities have been provided by SHARCNET (www.sharcnet.ca), Compute Canada (www.computecanada.ca) and the Department of Physics, Faculty of Science, Kasetsart University.

## Table of Content

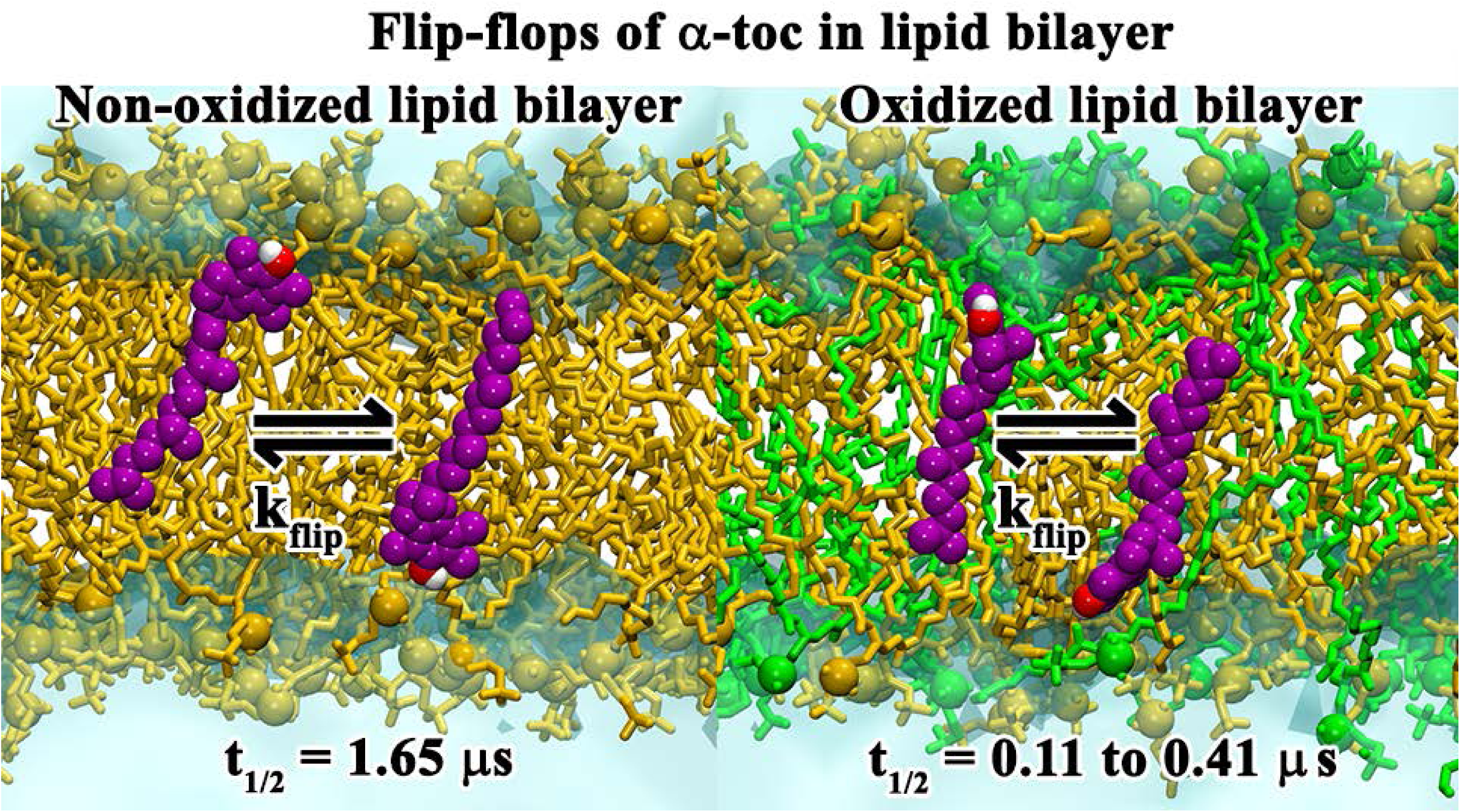

